# An ECM scaffold combined with a compliant 3D printed spring-shaped reinforcement for cartilage engineering applications

**DOI:** 10.1101/2024.01.31.575642

**Authors:** William Solórzano Requejo, Blanca Limones Ahijón, Carlota Corchado, Javier Llorca, Andrés Díaz Lantada, Jennifer Patterson, Pedro J. Díaz-Payno

## Abstract

Articular cartilage is a soft tissue lining the ends of the long bones in our joints. Even minor lesions in articular cartilage (AC) can cause underlying bone damage creating an osteochondral (OC) defect. OC defects can cause pain, impaired mobility and can develop osteoarthritis (OA). OA is a disease that affects nearly 10% of the population worldwide, and represents a significant economic burden to patients and society. While significant progress has been made in this field, realising an efficacious therapeutic option for unresolved OA remains elusive and is considered one of the greatest challenges in the field of orthopaedic regenerative medicine. Therefore, there is a societal need to develop new strategies for AC regeneration. In recent years there has been increased interest in the use of tissue-specific aligned porous freeze-dried extracellular matrix (ECM) scaffolds as an off-the-shelf approach for AC repair, as they allow for cell infiltration, provide biological cues to direct target-tissue repair and permit aligned tissue deposition, desired in AC repair. However, most ECM-scaffolds lack the appropriate mechanical properties to withstand the loads passing through the joint. One solution to this problem is to reinforce the ECM with a stiffer framework made of synthetic materials, such as polylactic acid (PLA). Such framework can be 3D printed to produce anatomically accurate implants, attractive in personalized medicine. However, typical 3D prints are static, their design is not optimized for soft-hard interfaces (OC interface), and they may not adapt to the cyclic loading passing through our joints, thus risking implant failure. To tackle this limitation, more compliant or dynamic designs can be printed, such as coil-shaped structures. Thus, in this study we use finite element modelling to create different designs including single triple, single quadruple, double triple and double quadruple helix and prototype them in PLA. The optimal design is combined with an ECM slurry. Briefly, the ECM slurry is combined with the PLA coil and freeze-casted under directional freezing prior to freeze-drying the samples to obtain an off-the-shelf scaffold with a dynamic reinforcement. The scaffold will be combined with mesenchymal stem cells (MSCs) to investigate the chondrogenic potential of such metamaterial. The double helix has a higher stiffness modulus than the single helix and the quadruple helix a higher stiffness modulus than the triple helix. The single helixes have a better recovery after compression, while the doble helixes have a higher plastic deformation under compression. The directional freeze-casting results in ECM scaffolds (either alone or PLA reinforced) containing a tailored microarchitecture mimicking aspects of native AC. To conclude, it was possible to design and simulate stiffness of different coil-shaped reinforcements and 3D print the PLA prototypes without support material. We were able to isolate and incorporate ECM into the coil structure and produce dynamic scaffolds that have the potential to be used in cartilage tissue engineering.

**Graphical abstract:** 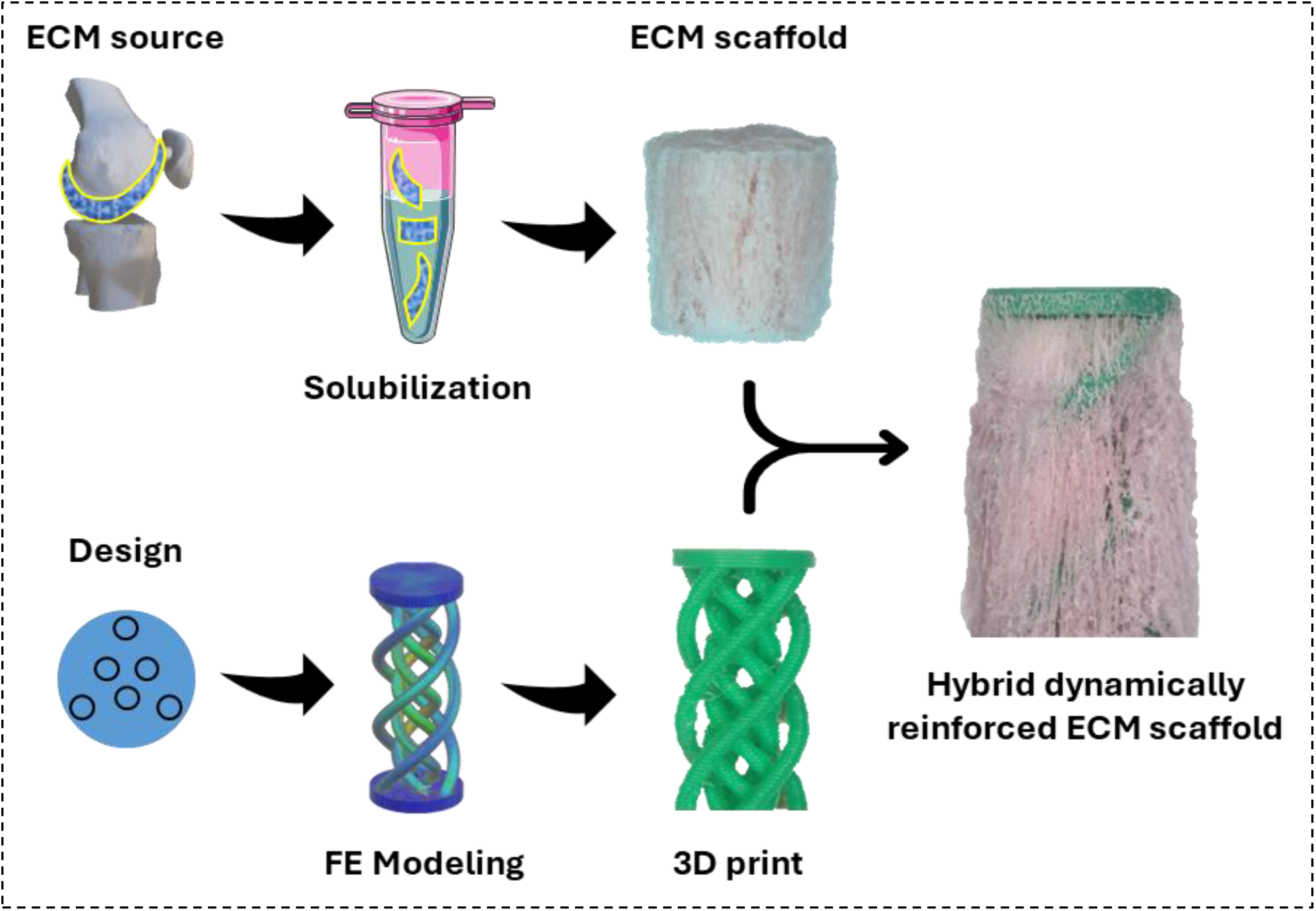

## 1. Introduction

Articular cartilage (AC) is a soft tissue lining the ends of the bones in our joints. Even minor lesions in AC can cause underlying bone damage creating an osteochondral (OC) defect. OC defects cause pain, impaired mobility and can develop to Osteoarthritis (OA). OA is the most common form of arthritis, affecting nearly 10% of the population worldwide[1], and it is a serious disease representing a significant economic burden to patients and society. In Europe, the cost of OA per patient is estimated to exceed € 10,000 per year [2]. At present, the treatment options for OA are limited to surgical replacement of the diseased joint with a prosthesis. While this procedure is well established, it is not without its limitations, and failures are not uncommon. In addition, joint replacement prostheses have a finite lifespan, making them unsuitable for the growing population of younger and more active patients requiring treatment for OA. While significant progress has been made in this field, realising an efficacious therapeutic option for the treatment of unresolved OA remains elusive and is considered to be one of the greatest challenges in the field of orthopaedic regenerative medicine[3]. One of the critical problems when repairing OC defects is the poor repair of the AC with a low-quality scar tissue, lacking the native aligned collagen microarchitecture[4]. Therefore, there is a societal need to develop new strategies for AC regeneration.

In the recent years there has been increased interest in the use of porous freeze-dried cartilage ECM derived scaffolds as an off-the-shelf approach for AC tissue repair, as they have a porous structure that allow for cell infiltration and provides the appropriate biological cues to direct target-tissue repair[5]. However, most of the freeze-dried ECM scaffolds focused on cartilage repair are fabricated with a homogenous pore structure that lacks a tailored pore microarchitecture to direct the alignment of tissue deposition, desired in AC repair. Recently, directional freeze-drying has been used to generate aligned pores that can allow for aligned tissue deposition. Furthermore, ECM-derived biomaterials lack the appropriate mechanical properties to withstand the loads passing through our joints [6], which may be hampering functional tissue repair. One solution to this problem is to reinforce the ECM with a stiffer framework made of synthetic materials, such as PLA [7]. Such framework can be 3D printed to produce anatomically accurate implants[8], attractive in personalized medicine. However, typical 3D printed structures are static, their design is not optimized for soft-hard interfaces (*e.g*. ECM-PLA interface), and they may not adapt to the cyclic loading passing through our joints, thus risking implant failure. To tackle this limitation, more compliant or dynamic designs can be printed, such as coil/spring-shaped structures[9]. This represents an improvement of the current implant design.

In order to succeed, two objectives are proposed: (i) The first objective consists of 3D printing a coil-shaped PLA structure with mechanical properties mimicking articular cartilage tissue. AC ECM-derived hydrogels and scaffolds that have been shown to support chondrogenesis of bone marrow derived stromal cells (MSCs) typically have poor mechanical properties[10], making them unsuitable for immediate load-bearing applications. One solution has been focused on reinforcing such ECM with synthetic 3D printed structures. However, such 3D printed constructs are static, and may not be well-suited to endure the cyclic loadings passing through the joints. We aim to design, and 3D print (using fused deposition modelling of FDM) a fully degradable coil-shaped structure made of medical grade PLA that is both immediately load-bearing and can adapt to the dynamic environment of the knee joint. (ii) The second objective focuses on the fabrication of a scaffold by directionally freeze-drying the PLA-reinforced ECM construct. The fabrication of ECM scaffolds with aligned pores has been successful through the directional freeze-casting of an ECM slurry before freeze-drying. Porcine AC ECM will be used as it closely mimics the human extracellular matrix [11]. We aim to adapt the freeze-casting protocol to the addition of the PLA structure and to test different % of AC ECM (0.5-3 %) in order to obtain a desired pore morphology (alignment and size) in the reinforced scaffold. Therefore, this phase will be involved in the characterization of the internal pore morphology (pore alignment and pore size).

## 2. Results and discussion

While significant progress has been made in the field of regenerative medicine, realizing an efficacious therapeutic option for the treatment of unresolved OA remains elusive and is considered one of the greatest challenges in the field of orthopaedic regenerative medicine[3]. A critical problem when repairing OC defects is the poor repair of the AC with a low-quality scar tissue, lacking the native aligned collagen microarchitecture[4]. Therefore, there is a societal need to develop new strategies for AC regeneration. Here, we have introduced a compliant or dynamic structural design for a complex reinforcement that can be 3D printed and combined with an ECM-derived scaffold.

### 2.1. Linear compression simulation shows the highest stress occurring in the inner face of the internal helix

It was possible to design the structures with Siemens NX. Briefly, a flat cylinder was used as base from which a coil thread emerged until reaching a second flat cylinder that was used as top. Different amounts of coil numbers were explored: 3 coils also referred as triple helix or 4 coils, quadruple helix. The structures were also designed to contain more than one group of helix to have either a single or a double helix.

Simulation of a linear compression test with 1N, 10 N and 50 N of the double triple helix and the double quadruple helix was performed with Siemens NX. The simulation demonstrated that the double triple helix could reach higher stress than the double quadruple helix (Figure 1A), reaching 12.7 MPa and 9.2 MPa, respectively. We hypothesize that the triple helix can be compressed more with the same amount of forced applied, and therefore the coil suffers a higher deformation and stress. In addition, the highest stress it is observed to occur in the inner face of the internal helix for both samples.

**Figure 1.**
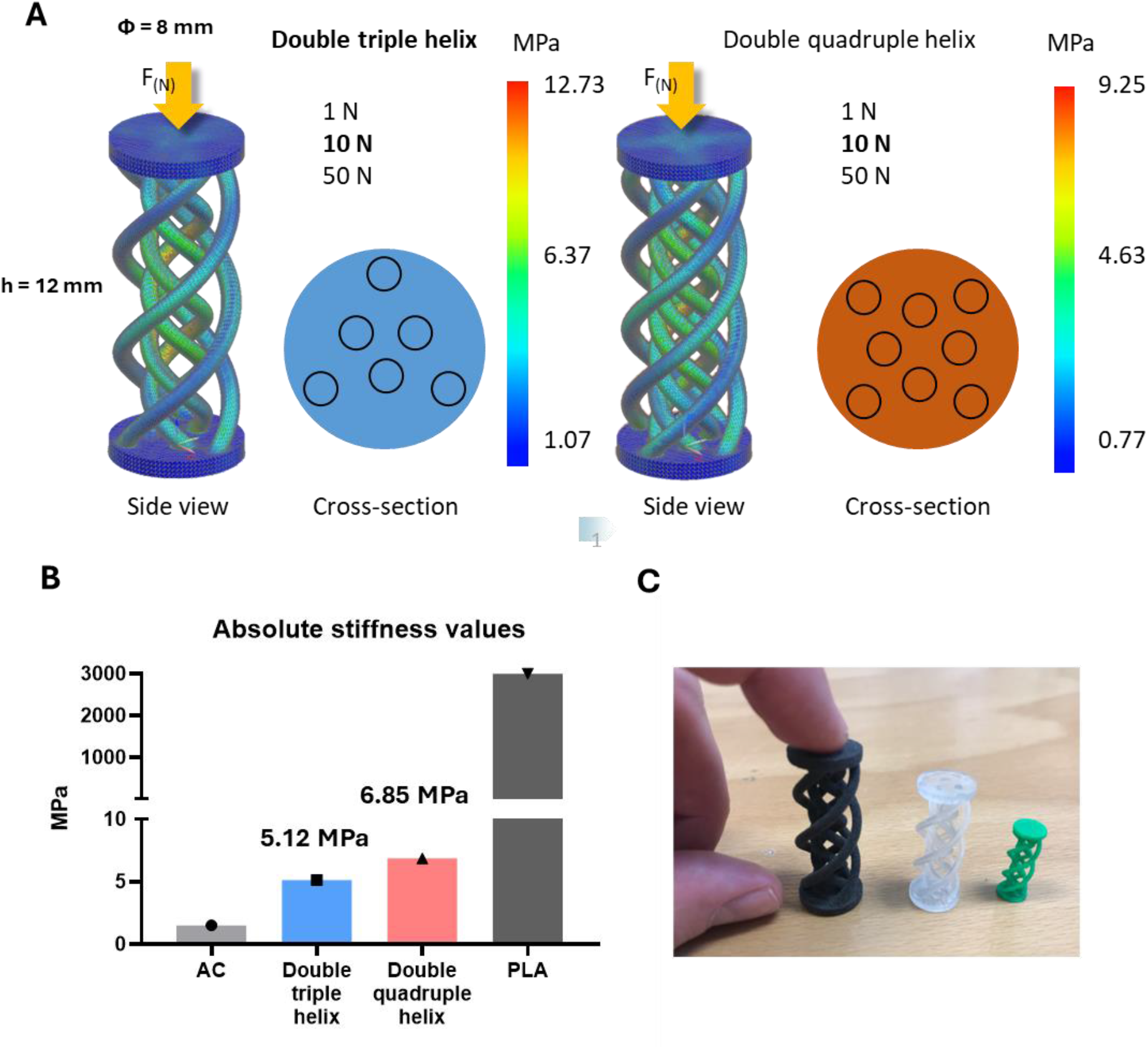
(A) Simulation of a linear compression test of the coil-shaped scaffolds for both double triple helix (blue) and double quadruple helix (orange). (B) Absolute stiffness values from simulation of linear compression test for both samples. Both AC and PLA compression modulus are included as comparison. (C) Double triple helix 3D printed with different materials and techniques to validate the printing feasibility of the structures: TPU (black), resin (translucid), green (PLA).

The theoretical stiffness values observed during the simulation when using standard parameters from PLA parameters as input for the material properties were 5.12 and 6.85 MPa for the double triple helix and double quadruple helix, respectively. Thus, the more helix the structure had, the stiffer the sample (Figure 1B). Different materials such as TPU, resin and PLA were used to 3D print the coil-shaped structures to test feasibility of the design (Figure 1C).

### 2.2. Coil-shaped structures can be 3D printed without support structures

Despite the overhanging nature of the coil shape, the structure could be 3D printed without the need for support material as seen in Figure 2. See supplementary video for more details on the printing process.

**Figure 2.**
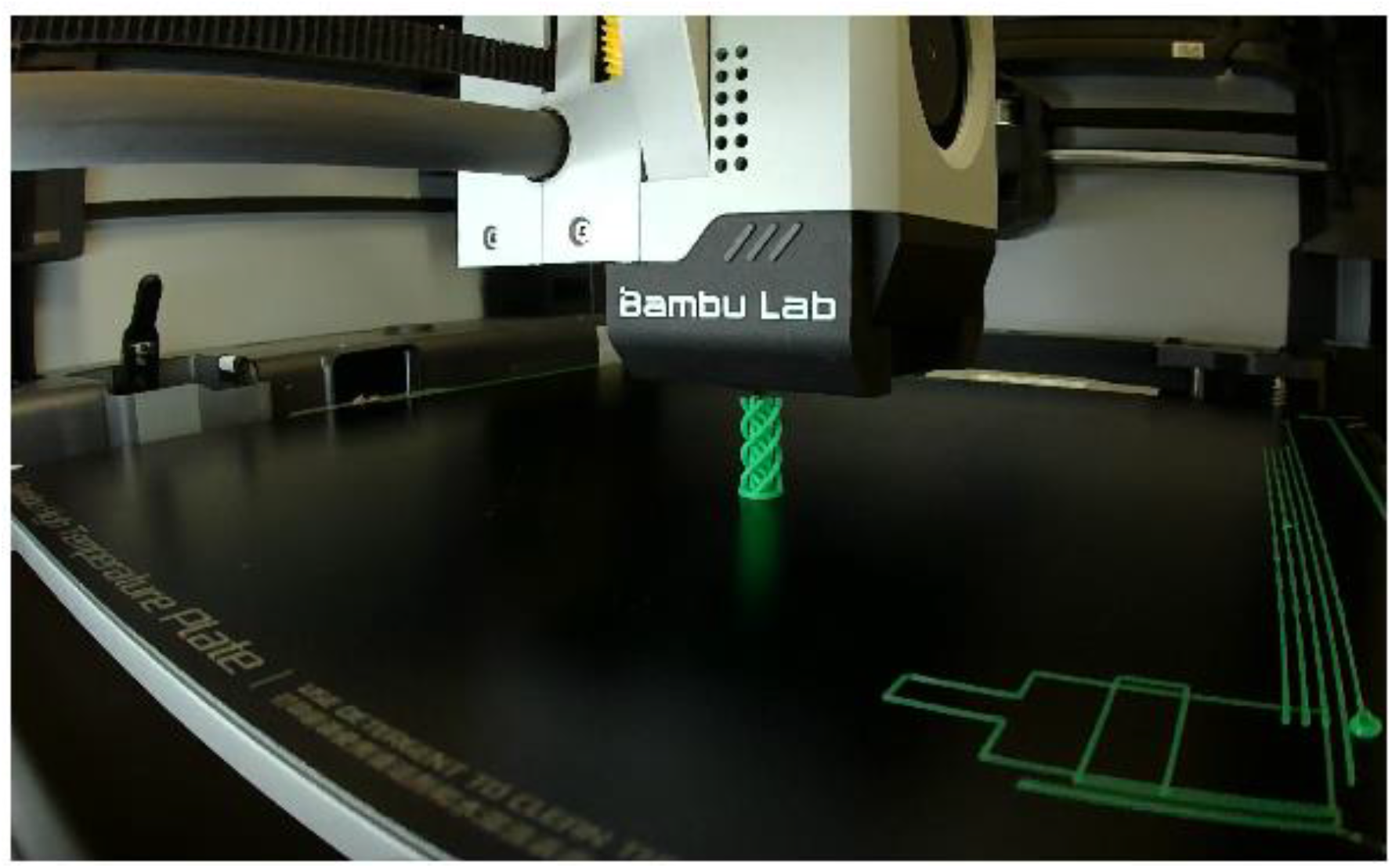
Photo in the middle of the 3D printing process of the coil-shape structures demonstrating the lack of supports needed to achieve the complex curved structures.

### 2.3. Scaling down of reinforcement dimensions and removal of internal helix lowers the mechanical properties and increases the recovery post-compression

We explored four different types of structures: single triple helix, single quadruple helix, double triple helix and double quadruple helix. The removal of the internal helix for the first two mentioned groups allowed to have a cylindrical hole that was hypothesized to facilitate the incorporation of the ECM derived material (Figure 3A). The STL file representative images capture the complex curvature and design of the four groups (Figure 3B).

**Figure 3.**
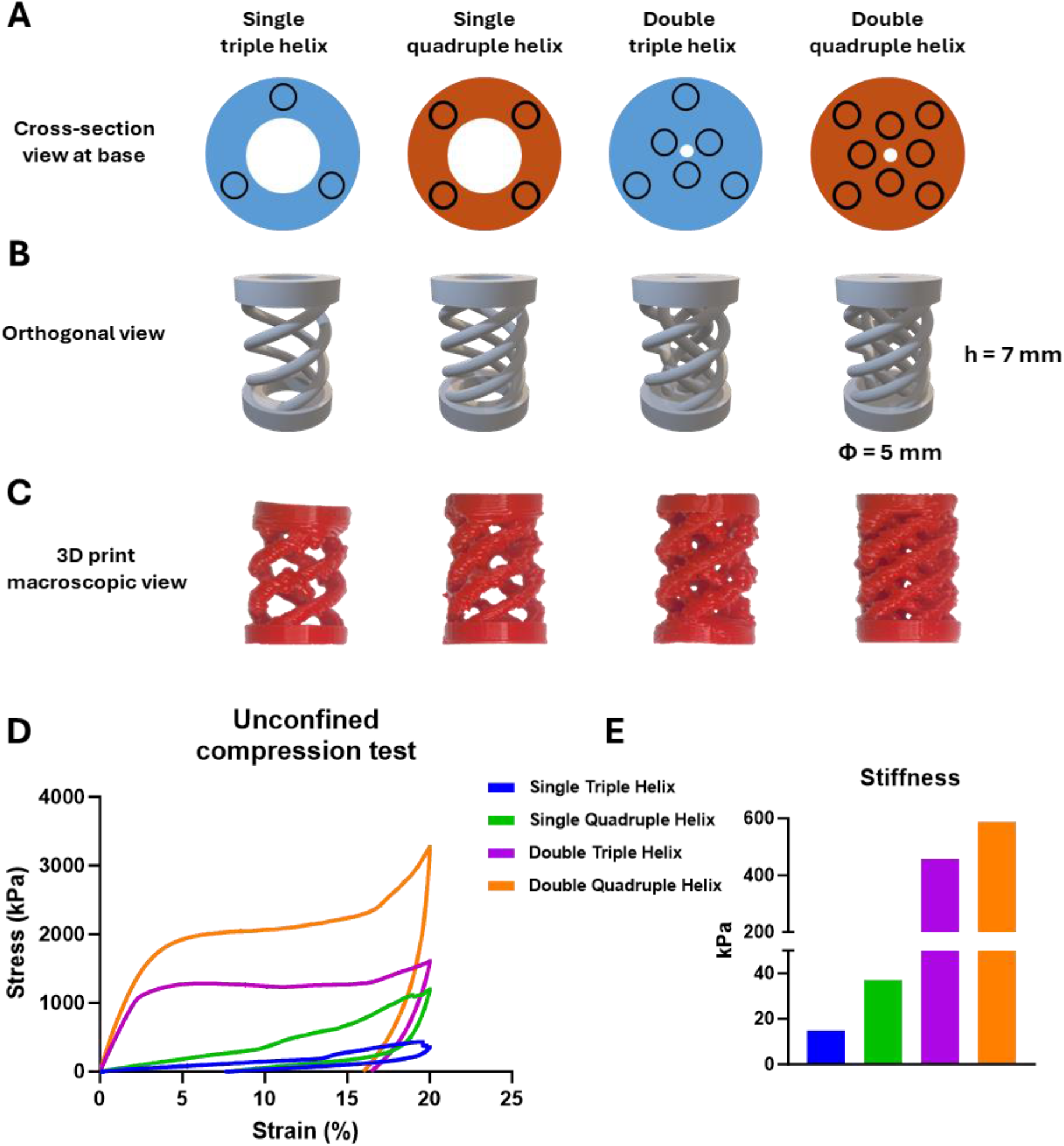
(A) Cross-section of the view from the base, illustrating the different number of coils and helix for the four samples. (B) STL file 3D representation for the 4 groups and (C) the macroscopic images of the 3D printed structures. (D) Stress-strain curve of the linear compression test and (E) compression Young’s modulus of the samples calculated from the first linear region of the stress-strain curve.

The designs were downsized for suitability during future cell culture studies in terms of ECM used, cells to seed and medium to feed the constructs. The structures were still able to be printed without any support material (Figure 3C).

Mechanical compression test was performed on the samples and it was observed in the stress-strain curve that the double helix groups suffered plastic deformation and were not able to recover, while the single helix groups did not have an apparent plastic deformation and they were able to recover (Figure 3D).

As simulated before, the highest stiffness corresponded to the double quadruple helix resulting in ∼600 kPa. The double triple helix had a young modulus of ∼400 kPa. The single quadruple helix had a young modulus of ∼40 kPa, while the single truple helix was ∼20 kPa.

### 2.4. Directional freeze-casting and annealing pre-freeze-drying results in an anisotropic pore structure with tailored pore size that is compatible with the addition of the coil-shaped PLA reinforcement

It was possible to isolate the extracellular matrix from the articular cartilage of porcine femoral joints. The matrix was solubilized as previously described before [10]. Briefly, the cartilage was biopsied from the femoral joint, treated with NaOH to remove the sGAGs, then digested with pepsine enzyme in acidic conditions to solubilize the tissue. The solubilized tissue was then salt extracted to concentrate the collagen fraction of the ECM. A schematic of the process can be found in the Supplementary information.

The solubilized ECM was used to fabricate freeze-dried scaffolds (Figure 4A). The scanning electron microscopy (SEM) imaging shown in Figure 4B demonstrated that the average pore diameter of the scaffolds was 70 µm, with a big distribution reaching up to 200 µm (Figure 4C). Orientation J plugin from Fiji software was used to characterize the alignment of the pores and demonstrated that the scaffolds had an anisotropic pore morphology distribution (Figures 4D, E).

**Figure 4.**
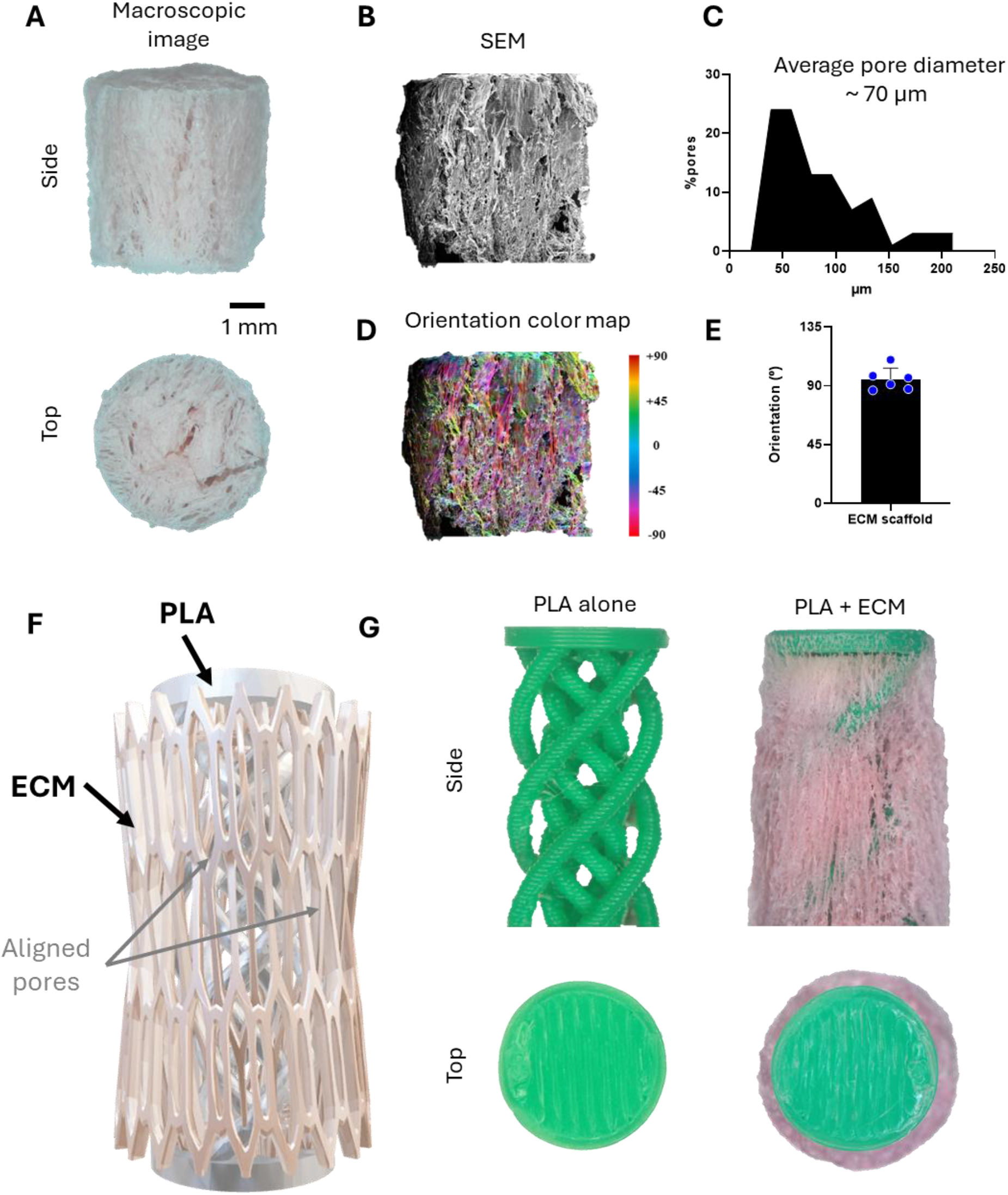
(A) Macroscopic images of the freeze-dried ECM scaffold. (B) SEM image of a cross-section of the scaffold. (C) Pore size analysis. (D) Orientation color map illustrating the anisotropy of the pore microarchitecture. (E) Quantification of the orientation. (F) Schematic of the combined PLA and ECM material into a hybrid scaffold. (G) Macroscopic images of the 3D printed PLA reinforcement (left) and the hybrid PLA-reinforced ECM scaffold (right).

Then we sought to combine the PLA coil-shaped reinforcement and the solubilized ECM. We demonstrated that this was possible, and that the orientation of the pores is observed to maintain a certain anisotropy. However, further research needs to be done to fully characterize the effects of the combination of both materials on the internal microarchitecture of the ECM scaffold (Figure 4F, G)

## Conclusion

Typical designs of 3D print reinforcements are static and may not adapt to the cyclic loading passing through our joints, thus risking implant failure. Here, we demonstrate that it was possible to design coil-shaped structures and 3D print them with PLA to generate dynamic reinforcement. We combined this reinforcement with ECM to fabricate off-the-shelf scaffolds that have a high potential for cartilage and osteochondral tissue engineering applications.

## Acknowledgements

This project has received funding from the European Union’s Horizon Europe research and innovation MSCA PF programme under grant agreement No. 101110000.

Vanesa Martínez – Mechanical characterization

Manuel Avella – SEM

Amalia San Román Suárez– Macroscopic, Sputtering pre-SEM

Carlota Largo Aramburu – Animalario idiPaz

## Notes

### Competing Interest Statement

The authors have declared no competing interest.

## References

[1] D.T. Felson, R.C. Lawrence, P.A. Dieppe, R. Hirsch, C.G. Helmick, Osteoarthritis: the diesease and its prevalaence and impact. In Felson DT, conference chair, Osteoarthritis: new insights. Part 1: The diesease and its risk factors, Ann Intern Med 133 (2000) 635–636.

[2] D.J. Hunter, D. Schofield, E. Callander, The individual and socioeconomic impact of osteoarthritis., Nat Rev Rheumatol 10 (2014) 1–5. 10.1038/nrrheum.2014.44.

[3] D.J. Hunter, S. Bierma-Zeinstra, Osteoarthritis, The Lancet 393 (2019) 1745–1759. 10.1016/S0140-6736(19)30417-9.

[4] M. Benders, Osteochondral defect repair: back to nature’s template, (n.d.).

[5] G.M. Cunniffe, P.J. Díaz-Payno, E.J. Sheehy, S.E. Critchley, H. V. Almeida, P. Pitacco, S.F. Carroll, O.R. Mahon, A. Dunne, T.J. Levingstone, C.J. Moran, R.T. Brady, F.J. O’Brien, P.A.J. Brama, D.J. Kelly, Tissue-specific extracellular matrix scaffolds for the regeneration of spatially complex musculoskeletal tissues, Biomaterials 188 (2019) 63–73. 10.1016/j.biomaterials.2018.09.044.

[6] N.E. Putra, K.G.N. Borg, P.J. Diaz-Payno, M.A. Leeflang, M. Klimopoulou, P. Taheri, J.M.C. Mol, L.E. Fratila-Apachitei, Z. Huan, J. Chang, J. Zhou, A.A. Zadpoor, Additive manufacturing of bioactive and biodegradable porous iron-akermanite composites for bone regeneration, Acta Biomater 148 (2022) 355–373. 10.1016/j.actbio.2022.06.009.

[7] M.T. Rodrigues, S.J. Lee, M.E. Gomes, R.L. Reis, A. Atala, J.J. Yoo, Bilayered constructs aimed at osteochondral strategies: the influence of medium supplements in the osteogenic and chondrogenic differentiation of amniotic fluid-derived stem cells., Acta Biomater 8 (2012) 2795– 806. 10.1016/j.actbio.2012.04.013.

[8] A.C. Daly, G.M. Cunniffe, B.N. Sathy, O. Jeon, E. Alsberg, D.J. Kelly, 3D Bioprinting of Developmentally Inspired Templates for Whole Bone Organ Engineering, Adv Healthc Mater 5 (2016) 2353–2362. 10.1002/adhm.201600182.

[9] H.M.A. Kolken, A.F. Garcia, A. du Plessis, C. Rans, M.J. Mirzaali, A.A. Zadpoor, Fatigue performance of auxetic meta-biomaterials, Acta Biomater 126 (2021) 511–523. 10.1016/j.actbio.2021.03.015.

[10] D.C. Browe, P.J. Díaz-Payno, F.E. Freeman, R. Schipani, R. Burdis, D.P. Ahern, J.M. Nulty, S. Guler, L.D. Randall, C.T. Buckley, P.A.J. Brama, D.J. Kelly, Bilayered extracellular matrix derived scaffolds with anisotropic pore architecture guide tissue organization during osteochondral defect repair, Acta Biomater 143 (2022) 266–281. 10.1016/j.actbio.2022.03.009.

[11] B. Liu, S. Zhang, W. Wang, Z. Yun, L. Lv, M. Chai, Z. Wu, Y. Zhu, J. Ma, L. Leng, Matrisome Provides a Supportive Microenvironment for Skin Functions of Diverse Species, ACS Biomater Sci Eng 6 (2020) 5720–5733. 10.1021/acsbiomaterials.0c00479.

