## Supplementary information for "An ECM scaffold combined with a compliant 3D printed spring-shaped reinforcement for cartilage engineering applications"

[1]ETSI Industriales, Polytechnic University of Madrid, Madrid, Spain
[2]Biomaterials and Regenerative Medicine group, IMDEA Materials Institute, Getafe, Madrid, Spain
[3]Bio/Chemo/Mechanics of Materials group, IMDEA Materials Institute, Getafe, Madrid, Spain
[4]Department of Materials Science, Polytechnic University of Madrid, Madrid, Spain
*Corresponding authors

### Scaffold fabrication

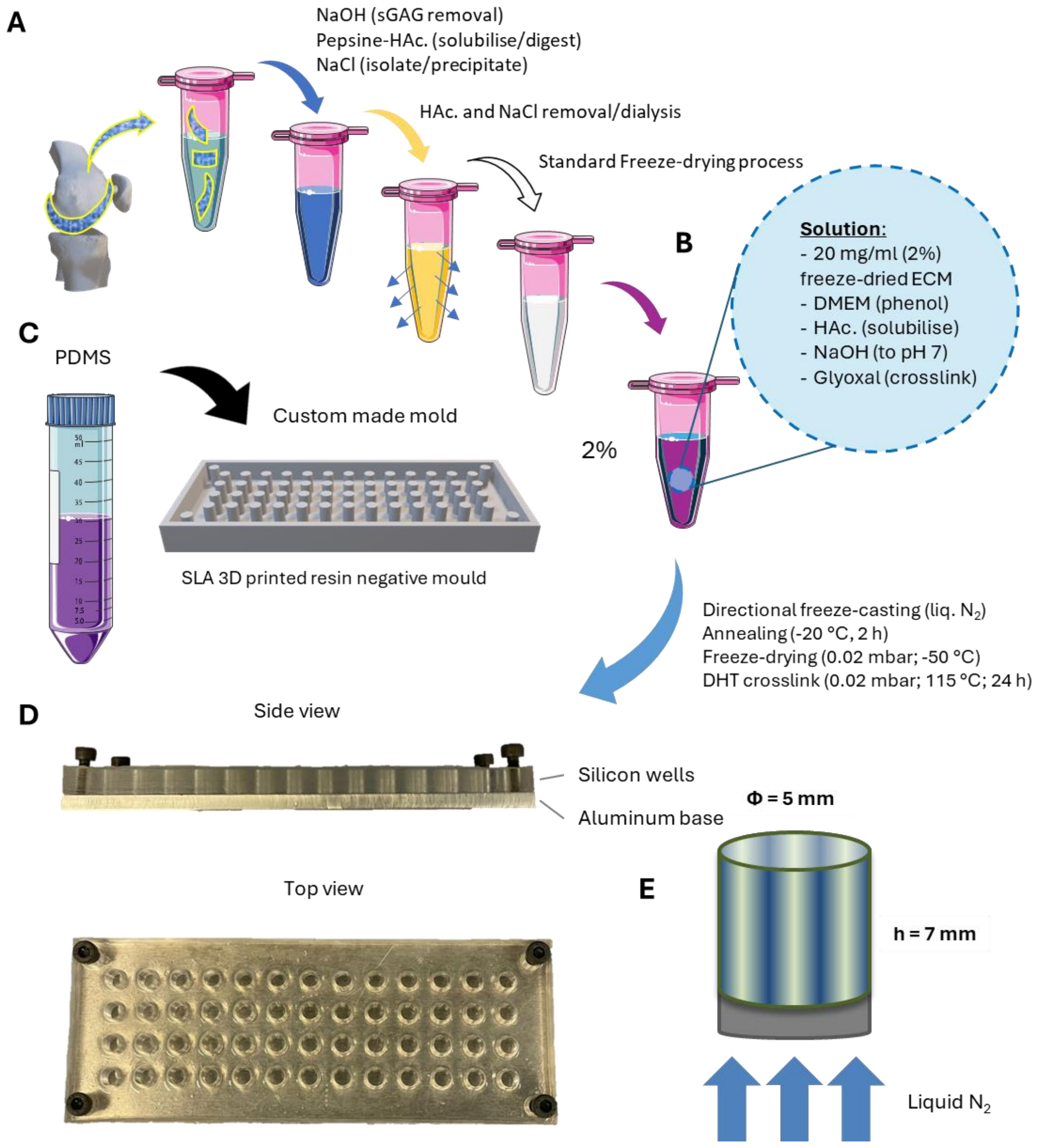
